# Reappraising the human mitochondrial DNA recombination dogma

**DOI:** 10.1101/304535

**Authors:** Simόn Perera, Amanda Ramos, Luis Alvarez, Débora Jurado, Maria Guardiola, Manuela Lima, Maria Pilar Aluja, Cristina Santos

## Abstract

With the “mitochondrial Eve” theory proposed by Rebecca Cann in the eighties, human mitochondrial DNA (mtDNA) has been used as a tool in studying human variation and evolution. Although the existence of recombination in human mtDNA has been previously advocated, studies dealing with human variation and evolution have assumed that human mtDNA does not recombine and should be considered as pathological or very infrequent. Using both direct and indirect approaches, we provide consistent evidence of mtDNA recombination in humans. We applied the single molecule PCR procedure to directly test for recombination in multiheteroplasmic individuals without any overt pathology. Moreover, we searched for past recombination events in the whole mitochondrial genomes of more than 15,000 individuals. Results from our study update and expand both the seminal indirect findings and the scarce direct evidence observed to date, paving the way for the definitive rejection of the non-recombination dogma for human mtDNA. Acknowledgment of recombination as a frequent event in mtDNA will require the description of the population recombination rate(s) and to apply it to past and future studies involving mtDNA. MtDNA recombination affects our knowledge of human evolutionary history, regarding haplogroup divergence times, as well as the time to the mitochondrial most recent common ancestor. Finally, mtDNA recombination will have a substantial impact on our understanding of the etiology and transmission of mitochondrial diseases.

## Introduction

The value of human mitochondrial DNA (mtDNA) as a major tool for evolutionary studies lies in its small size, the high number of copies per cell, a mutation rate higher than nuclear DNA, its maternal inheritance and the absence of recombination. Although it is currently thought that mtDNA recombination does not occur, evidences accumulated during the last twenty years have challenged this “dogma”. In the 1990s when the enzymatic machinery of recombination was identified in mitochondria (1), the absence of recombination was first questioned. Later, intramolecular recombination was observed in human cell cultures (2) and recombination intermediates were detected (3, 4). Despite these arguments, the absence of mtDNA recombination in humans has long been assumed. Currently, there is consistent evidence that, in humans, the paternal transmission of mtDNA is a very infrequent event (5), in fact paternal mtDNA appears to be eliminated at fecundation and until now, a single case has been reported (6). Interestingly, the first direct evidence of human mtDNA recombination was obtained over 10 years ago by Kraytsberg et al. (7) in the muscle tissue of this previously reported case, where paternal and maternal mtDNAs were mixed (6).

MtDNA copies present in a given individual may be nearly identical, which limits the observation of recombination. Notwithstanding, different mtDNA molecules can coexist in the same individual, a condition known as heteroplasmy, that nowadays is considered to be universal. In this sense, multiheteroplasmic individuals, in which recombination could potentially create new allele combinations (mitotypes), would be essential to empirically detect recombination in mtDNA. Using this special feature of mtDNA, Zsurka et al. (8) analysed the distribution of allelic combinations in patients with neurological disorders and multiple mtDNA heteroplasmies, finding a mixture of the four possible mitotypes, which the authors considered a hallmark of recombination.

Besides the empirical evidences of mtDNA recombination, several indirect methods have been developed and applied to infer past recombination events in mtDNA (9). Eyre-Walker et al. (10), among others, indirectly found the existence of recombination, but their results were later questioned, and conclusions reported as unreliable (11). Most recently, White et al. (12) concluded that often used indirect tests, such as Max *χ*^2^ test, are unsuitable to detect recombination in mtDNA and that the neighbour-similarity score (NSS) (13) is the most reliable available method to test for recombination in mtDNA. Moreover, White et al. (14) showed that mtDNA recombination has only been tested in small datasets, well below the now available compiled information of mtDNA sequences.

Here we test for the presence of recombination in human mtDNA and provide consistent evidence for widespread recombination using both direct and indirect approaches.

## Materials and Methods

### 1. Direct evidence of recombination

#### Sample collection and DNA extraction

From a sample of 101 individuals previously analysed by Ramos et al. (15), two double-heteroplasmic individuals were identified and selected for further described analysis, Z141 and Z176. Appropriate informed consent and the known birth places of maternal ancestors (up to the third generation) had been obtained under confidentiality (15). The present study was approved by the ethics committee of the Universitat Autònoma de Barcelona. Ramos et al. (15) detected and validated the heteroplasmic positions in the whole mtDNA sequences. Individual Z141 is heteroplasmic in positions 189 and 15496 [according to the revised Cambridge reference sequence (rCRS) (16)], with 85.1 % <u>A</u> / 14.9 % G in position 189 (ancestral variant underlined), and 86.2 % G / 13.8 % <u>A</u> in position 15496. Individual Z176 is heteroplasmic in positions 8307 and 15908 [according to the rCRS (16)], with 66.3 % <u>A</u> / 33.7 % G in position 8307 (ancestral variant underlined), and 73.0 % <u>T</u> / 27.0 % C in position 15908 (15).

#### Single-molecule PCR

To test the selected individuals for recombination, the single-molecule PCR (smPCR) procedure was followed (17) (Supplemental Figure S1). In brief, we serially diluted DNA to reach the highest dilution containing one or a few amplifiable DNA templates while the following dilution should not contain any DNA templates. After adjusting for the Poisson distribution, we worked with a 0.3 fraction of positive events, which corresponds to 0.36 real templates and 0.05 multiples in each positive reaction.

To perform the smPCR in Z141 we selected the primers 14898for (5′- tagccatgcactactcaccaga-3′) and 1677rev (5′-gtttagctcagagcggtcaagt-3′) from Ramos et al. (18), allowing for the amplification of a ~3350bp region of the mtDNA containing both heteroplasmic positions. Concerning Z176, primers 7713for (5′- tcctaacactcacaacaaaac-3′) and 16281rev (5′-gttggtatcctagtgggtgag-3′) were used (18) to amplify ~8568bp.

The smPCR mix for each sample consisted in 10 pM of each primer, 2.5 μM of dNTP mix, *10x LA PCR Buffer II* (*Mg2+plus*), 0.5 U of LA Taq polymerase (Takara Bio Europe, Saint-Germain-en-Laye, France), and 1 μL of sample (at the concentration established from the serial dilution procedure). The smPCR programme, performed in an S1000 thermocycler (Bio-Rad, Hercules, USA) comprised an initialization step of 1 min at 94 °C, 30 PCR cycles (denaturation: 30 s at 94 °C, annealing: 60 s at 60 °C, elongation: 3 min for Z176 / 6min for Z141 at 72 °C) and a final step of 10 min at 72 °C.

#### Nested PCR

Positive reactions from the smPCR were subjected to nested PCR. Independent amplifications were performed for each of the heteroplasmic positions (Z141-position 189: primers L2-16485 5′-gaactgtatccgacatctgg-3′(19) and H2-481 5′- gattagtagtatgggagtgg-3′(19); Z141-position 15496: primers 15416for 5′- tacacaatcaaagacgccctc-3′(18) and 15825rev 5′-gtgaagtatagtacggatgct-3′(18); Z176- position 8307: primers 8196for 5′-acagtttcatgcccatcgtc-3′(20) and 8600rev 5′- agaatgatcagtactgcggcg-3′(18); Z176-position 15908: 15721for 5′-ttgactcctagccgcagac-3′(21) and 16042rev 5′-ctgcttccccatgaaagaacag-3′(21).

The nested PCR mix consisted in 50 pM of each primer, dNTP mix (100 μM), *10x NH_4_ Reaction Buffer*, MgCl_2_ (50 mM), 0.5 U of BIOTAQ polymerase (Bioline, Taunton, USA), and 1 μL of smPCR product. The PCR programme consisted in an initialization step of 5 min at 95 °C, 35 PCR cycles (1 min at 94 °C, 40 s at 55 °C, 1 min at 72 °C) and a final step of 5 min at 72 °C.

The amplification of each sample was verified through electrophoresis with 1.5 % agarose gel and 0.1 % ethidium bromide. To avoid false negatives due to the low DNA concentration after the smPCR this step was performed after the nested PCR.

#### Sequencing

Nested PCR products were sequenced to confirm the single-molecule status of their templates (a single base fluorescence signal can be observed in the position studied) and to establish the specific nucleotidic base of each mitochondrial molecule to allow the determination of their mitotype. Sequencing reactions were performed using the BigDye Terminator v3.1 Cycle Sequencing Kit. Sequencing products were purified through ethanol/EDTA precipitation, and reactions were obtained with an Applied Biosystems 3130XL sequencer (Servei de Genòmica, Universitat Autònoma de Barcelona).

### 2. Indirect evidence of recombination

#### Sequence gathering and alignment, haplogroup assignment and grouping

We extracted 15153 complete mitochondrial genomes from PhyloTree (build 16) (22) and assigned each sequence to its haplogroup (Hg), according to the PhyloTree nomenclature, using the HaploFind software (23) (Supplemental Figure S2). Only the highest-quality sequences were considered for analysis, according to their annotation in PhyloTree. Sequences previously determined as false recombinants, sequences obtained from articles including erroneous sequences, incomplete sequences and sequences inferred from haplotypes were not considered for the analysis.

Sequences were clustered in the monophyletic groups relevant for their posterior analysis, ranging from single-sequence subhaplogroups to the global set of sequences (see Suppementary data). The largest sets of sequences were divided in smaller sets, and sequences in each subset were aligned with the Muscle algorithm (24).

We have tested for the presence of recombination in: A) the entire set of sequences; B) macrohaplogroups (Mhgs) M, N*, R*, R0, and U; and C) haplogroups with more than 100 sequences. Asterisked Mhgs represent the sets of sequences belonging to one macrohaplogroup and not belonging to other Mhgs nested within it.

The whole set of sequences (Test A) was divided in 29 different subsets (Table 1; Supplemental Figure S2), with macrohaplogroup relative frequencies approximately corresponding to that of the whole analysed mtDNA sequence set. Macrohaplogroups M, N*, R*, R0, and U (Test B), were split in a total of 23 random subsets of sequences (Suppementary data). Thirty-six haplogroup-defined subsets were analysed (Test C) (Suppementary data).

**Table 1.** **Percentage of tested subsets positive for recombination.**Ranges correspond to the most and least conservative methods to consider subsets as positive for recombination. NSS: neighbour sequence similarity.

|  | % subsets positive for recombination |  |
| --- | --- | --- |
| | NSS | Max $\chi^2$ |
| **Global** | 86-95 | 3-7 |
| **Mhgs** | 60-90 | 4-13 |
| **Hgs** | 58-87 | 2-9 |

#### Presence of recombination

Two tests have been used to test for recombination: the neighbour sequence similarity (NSS) test (13) and the maximum χ^2^ (Max χ^2^) test (25, 26). The NSS test analyses the distribution of sites whose history may include recurrent or convergent mutation or recombination, whether the Max χ^2^ test compares the arrangement of segregating sites at either side of a putative crossover break point (12). All tests have been performed with the software package PhiPack (27).

Before analysing for recombination, polycytosines and their flanking regions (CRS positions 298-320 and 16179-16198) were deleted for the sake of alignment quality, as well as all positions containing at least one gap in the alignment.

Only the highest-quality sequences were considered for analysis, according to their annotation in PhyloTree. Sequences previously determined as false recombinants, sequences obtained from articles including erroneous sequences, incomplete sequences and sequences inferred from haplotypes were not considered for the analysis.

#### Statistical analysis

In order to correct for statistic type I errors in null hypothesis rejection, we used Benjamini-Hochberg’s false discovery rate (FDR) controlling procedure (28). The percentage of recombination in the tested subsets was assessed through two methods. In the most conservative one, only subsets with FDR = 0.05 were considered, the final quantity of positive subsets was corrected by this FDR, and numbers of valid subsets were truncated. In the least conservative method, the probability of each subset to be positive for recombination was calculated according to pre-defined thresholds of FDR = 05, 0.10, 0.25 or 0.50, each subset’s probability was corrected according to its corresponding FDR value, and numbers of valid subsets were rounded.

## Results and Discussion

### 1.Direct evidence of recombination

We tested for mtDNA recombination in two double-heteroplasmic individuals without any overt pathology previously identified by Ramos et al. (15): individual Z141 presenting heteroplasmy in positions m.189A>G and m.15496G>A, and individual Z176 presenting heteroplasmy in positions m.8307A>G and m.15908T>C. SmPCR, a procedure which successfully detects mtDNA recombination and avoids PCR artifacts (17), was used to detect the mitotypes in each individual. We successfully amplified and sequenced 78 single molecule-derived replicas from Z141 and 97 from Z176. In both cases all possible mitotypes were identified for both samples, evidencing the possible existence of mitochondrial recombinant products in the two healthy individuals (Figure 1). Apart from the mtDNA mutations in heteroplasmy analysed for the present study no other mutations were detected.

The heteroplasmic positions of Z141 appear in the control region (189 <u>A</u>/g, ancestral variant underlined) and cytochrome b (15496 G/<u>a</u>). The AG and GA mitotypes can be explained by the mutations which cause the heteroplasmies from an ancestral mitotype AA (29). Regarding the appearance of the minoritary GG mitotype, one possibility is for a third mutation to have occurred in each of the already mutant mitotypes, however this scenario implies recurrent mutations and although 189 is a hotspot position [45 hits in the mtDNA phylogeny (22)], 15496 is a stable position [one hit in the mtDNA phylogeny and a probability of mutation of 0,00019 (15)], making the recurrence of this mutation highly unlikely. Thus, it is probable that the GG mitotype has originated through mitochondrial recombination. Furthermore, concerning the Z176 individual, the heteroplasmic mutations are both located in highly stable positions of tRNAs Lys and Thr (8307 <u>A</u>/g and 15908 <u>T</u>/c positions, respectively), presenting zero and one hit in the phylogeny and a probability of mutation of zero and 0,00019, respectively (15). Again, while the AC and GT mitotypes are explainable by direct mutation from the ancestral, still predominant, mitotype, the GC mitotype is very unlikely to have been originated through a third mutational event. Considering the data obtained altogether, only mtDNA recombination could be responsible of the mitotype distribution pattern observed in this individual (Figure 1).

**Figure 1.**
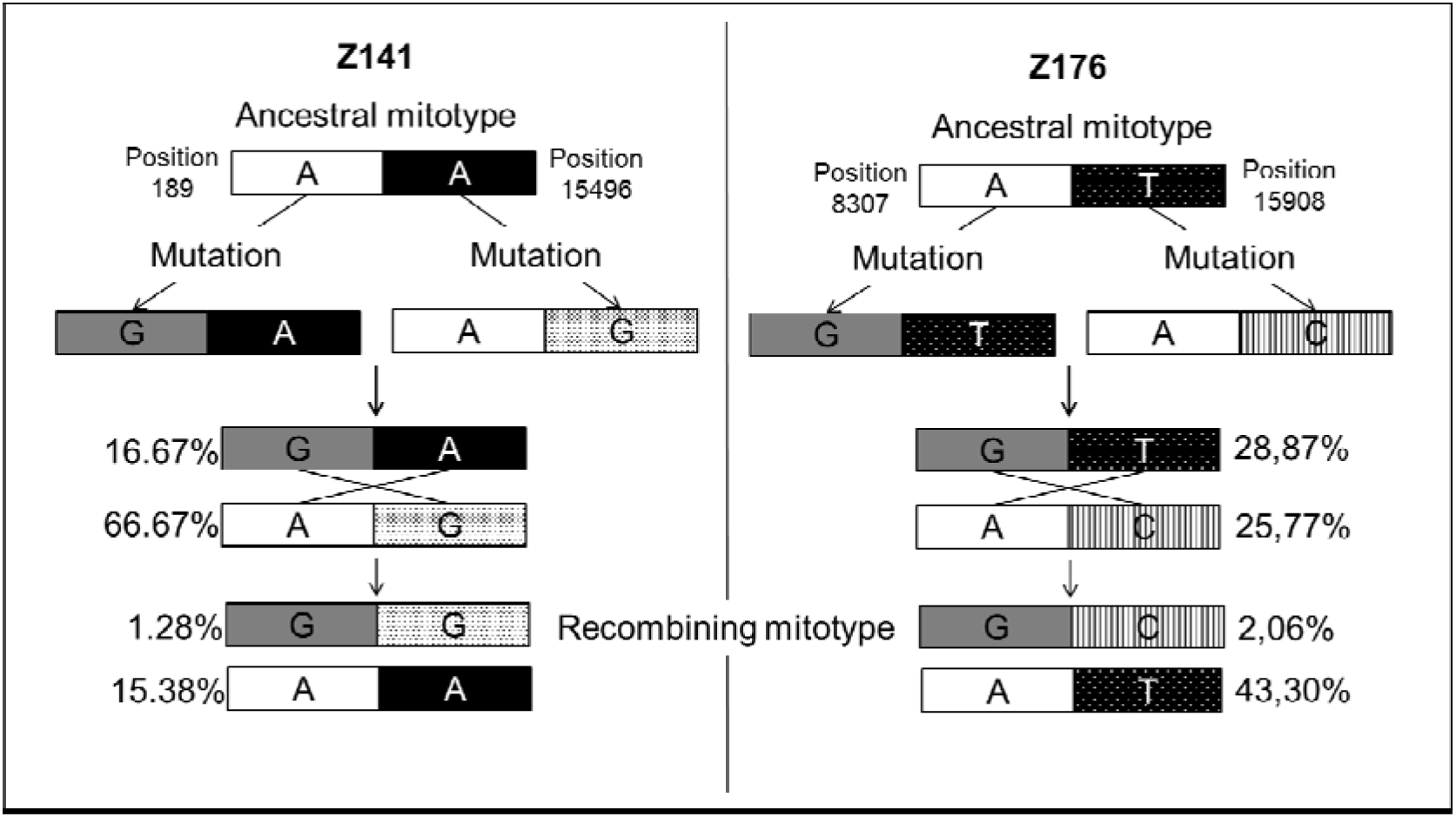
**Direct evidence of human mtDNA.**Frequency of mitotypes obtained by smPCR in the multiheteroplasmic individuals Z141 and Z176. Heteroplasmic positions analysed are also shown.

### 2.Indirect evidence of recombination

We tested for recombination in human mtDNA using 15153 complete human mitochondrial genomes from PhyloTree (build 16) (22) and assigned each sequence to its haplogroup (Hg). We analysed the existence of recombination in the global dataset, as well as in macro-Hg (Mhg) and Hg-defined sequence subsets, with the NSS test (13) and Max χ^2^ (25, 26) tests.

We found indirect evidence of past recombination events in 86-95 % of the global-sample subsets, 60-90 % of the Mhg-defined subsets, and 58-87 % of the Hg-defined subsets (Table 1; Suppementary data). Moreover, although the Max χ^2^ test is not suited to detect recombination in mtDNA (12), it found evidences of recombination in some subsets (Suppementary data). It is also noteworthy that recombination is observed in all Mhgs, as well as in their belonging Hgs.

Although the NSS test sometimes finds false positive (FP) recombination, in the worst-case scenario of low sequence diversity and a substitution rate correlation of 0.6 [as is the case for mtDNA (30)], the FP ratio is ~30 % (25), thus guaranteeing the presence of significant recombination in 45-64 % of our tested subsets. The evolution model can also affect the FP ratio of the applied tests. Particularly, NSS tends to overestimate recombination in *patchy-tachy* evolution, a model of substitution in which mutation rates vary across taxa for some parts of the sequences (9). Although evidence for *patchy-tachy* is null for humans, and contrasting for cercopithecids, correcting our results with simulation-based FP ratio values (9) in addition to the previous FP correction, we still find mtDNA recombination in ≥ 37-52 % of the subsets, thus unequivocally supporting the existence of recombination in human mtDNA.

The existence of recombination is particularly important because it may affect current demographic estimates derived from coalescent events, such as divergence times and patterns of population expansion. Our results point towards a widespread existence of recombination, affecting all haplogroups and thus having been present during at least all the recent evolutionary history of our species. Our work makes it necessary to describe the mtDNA population recombination rate(s) and apply it to past and future studies involving mtDNA. MtDNA recombination affects our knowledge of human evolutionary history, regarding haplogroup divergence time, as well as time to the mitochondrial most recent common ancestor. Finally, mtDNA recombination will have a substantial impact on our understanding of the etiology and transmission of mitochondrial diseases.

Overall, our results strengthen the evidence for mtDNA recombination: not only does mitochondrial recombination exist in isolated or diseased individuals, but it is widespread in human populations at all evolutionary levels. These results update and expand both the seminal indirect findings and the scarce direct evidence observed to date, paving the way for the definitive rejection of the non-recombination dogma for human mtDNA.

## Acknowledgements

This work was partially supported by Generalitat de Catalunya (2009SGR566, 2014SGR1420, 2017SGR1630). A postdoctoral fellowship (SFRH/BPD/105660/2015) (A.R.) was supported by Fundação para a Ciência e a Tecnologia (FCT).

## Author Contributions

The study was designed by S.P., A.R. and C.S.; Data analysis and interpretation was performed by S.P., A.R. and C.S.; Direct evidence study was performed by S.P, A.R., D.J. and M.G.; Indirect evidence study was performed by S.P., L.A and C.S.; MP.A., M.L. and C.S. funded the direct evidence studies; Sample acquisition, annotation and collection was performed by A.R., L.A., MP.A. and C.S.; S.P., A.R. and C.S. prepared and wrote the manuscript; All authors contributed revisions to the manuscript. Note that S.P. and A.R. contributed equally to this work.

